# Dynorphinergic neuroadaptations in the islands of Calleja: implications for alcohol use disorder

**DOI:** 10.64898/2026.06.05.730464

**Authors:** Anthony M. Cuozzo, Gaetan Lepreux, Daniel J. Reis, Gengze Wei, Brendan M. Walker

## Abstract

Dysregulation of the dynorphin (DYN) / kappa-opioid receptor (KOR) system is heavily implicated in symptoms of alcohol use disorder (AUD) including negative affective-like states that can drive maladaptive behavioral regulation. Substantial efforts have been made towards understanding the neurobiology of DYN / KOR dysregulation; however, the role of dynorphinergic islands of Calleja within the ventral striatum remain poorly understood. Presently, adult male Wistar rats were trained to self-administer 10% alcohol, exposed to either air or alcohol vapor for eight weeks, and alcohol self-administration and 22-kHz ultrasonic vocalizations (USVs) assessed during acute withdrawal. Subsequently, brains were extracted during acute withdrawal and DYN A-like immunoreactivity was measured in the ventral striatum. Alcohol vapor-exposed rats demonstrated increased alcohol consumption and 22-kHz USVs compared to air-exposed controls. Vapor-exposed rats additionally demonstrated increased DYN A-like immunoreactivity in the islands of Calleja. Moreover, the average DYN A neuron size positively correlated with the number of 22-kHz USVs in vapor exposed animals, but not in air-exposed controls. The present findings identify the islands of Calleja as a novel DYN-associated region that may be recruited during alcohol dependence with enhanced DYN plasticity in the islands of Calleja contributing to affective dysregulation in AUD and other neuropsychiatric conditions.

## Introduction

The Islands of Calleja (IC) are densely packed granule cell clusters [1] situated within the olfactory tubercle (OT). While earlier assessments focused on the OT within the context of olfaction preference [2], the IC are now emerging as a functionally distinct component of the ventral striatal complex. The IC receives dopaminergic (DA) input from the ventral tegmental area (VTA), as well as projections from cortical and other limbic regions [3, 4]. Given the IC integration within the ventral striatum, it is plausible that the IC contribute to the neural computations underlying reinforcement, behavioral flexibility, and emotional reactivity [5]. Recent evidence in mice has shown that the IC control grooming [6], a behavior associated with negative affective-like states when excessive. Moreover, IC activity is dysregulated following chronic restraint and contributes to depressive-like behaviors [6]. These findings suggest the IC may play a role in affective disturbances, serving as a target of stress-induced neuroadaptations or the emergence of negative emotional states.

The kappa-opioid receptor (KOR) is a important modulator of stress and dysphoria in both humans and animals [7]. KORs are inhibitory G-protein coupled receptors activated by the endogenous opioid peptide dynorphin (DYN) [8]. KORs are widely expressed throughout the brain, in regions implicated in emotion and reinforcement-related behaviors, including the VTA, nucleus accumbens (NAc), and amygdala [9]. Given its position in key regions of motivational circuitry, the KOR/DYN system has been established as a critical contributor to the maladaptive behavioral regulation that accompanies alcohol use disorder (AUD)[10].

KOR/DYN signaling plays an important role in the transition between controlled alcohol use to compulsive drinking in the context of AUD [10]. During withdrawal, increased KOR function contributes to the emergence of negative emotional states that produce enhanced alcohol-seeking behaviors [e.g., 11, 12-16] and KOR expression in VTA DA neurons play an important role in symptoms of AUD [14]. However, the precise anatomical substrates through which KOR/DYN signaling exerts its effects on affective states remain incompletely understood. The IC receive inputs from VTA DA neurons and are connected with important motivational and affective regions [1, 4], as such the IC represents a compelling target for further investigation given its role in affective behavior.

In the present study we evaluated DYN A-like immunoreactivity in the IC of Wistar rats made alcohol-dependent through a chronic intermittent alcohol vapor exposure regimen [11, 12]. Moreover, prior to the immunohistochemical processing, those animals were assessed during acute withdrawal for behaviors consistent with phenotypes of AUD.

## Methods

### Animals

Nine male Wistar rats approximately 70 days of age (bred from Charles River Laboratory breeding pairs) were housed in groups of two or three in this experiment. The environment was controlled for temperature and humidity in a reverse 12h light cycle with ad libitum food and water. Animals were handled for five consecutive days before the beginning of the study and animal care adhered to the National Research Council’s Guide for the Care and Use of Laboratory Animals with all procedures approved by the Washington State University and University of South Florida Institutional Animal Care and Use Committees.

### Operant self-administration of alcohol

Animals were trained to self-administer 10% ethanol (w/v) using a sweetener-fade procedure [see 17 for details]. Sessions were conducted in operant conditioning chambers (Med Associates, St. Albans, VT, USA) equipped with dual drinking wells (Behavioral Pharma, La Jolla, CA, USA) under an FR-1 schedule for 30-min. After acquisition and stabilization of responding, animals were assigned to either a vapor exposure group exposed to chronic intermittent ethanol vapor or an air-exposed control group. Following eight weeks of exposure, alcohol self-administration was assessed during acute withdrawal (6–8 h after vapor turned off).

### Chronic intermittent alcohol vapor

To induce dependence, male Wistar rats were subjected to chronic intermittent ethanol (CIE) vapor exposure (14 h on/10 h off daily) in custom built vapor chambers (La Jolla Alcohol Research, La Jolla, CA); for detailed methodology, see Williams et al., 2012. Ethanol vapor concentration was adjusted to maintain blood alcohol levels (BALs) between 175–250 mg/dl, monitored twice weekly via tail vein blood sampling and analyzed with an Analox GL5 (Analox USA, Atlanta, GA, USA) analyzer.

### Ultrasonic vocalization recordings

To assess negative affective-like behavior during acute withdrawal, 22-kHz ultrasonic vocalizations (USVs) were recorded 48 hours after the acute withdrawal self-administration session. Non-painful 60 psi air-puffs were used to elicit vocalizations [11, 18]. USVs were recorded using an ultrasonic microphone and analyzed with Avisoft software (Avisoft Bioacoustics, Berlin, Germany).

### Immunohistochemistry

DYN-A like immunoreactivity was assessed using a standard immunoperoxidase staining procedure as described previously [12]. Briefly, free-floating sections were incubated in anti-DYN A (1-17) primary antibody (1:1000; Peninsula Laboratories) for 20 hours, followed by incubation with a biotinylated anti-rabbit secondary antibody and development using an avidin-biotin complex and DAB substrate. Images were captured under uniform conditions and analyzed using ImageJ to quantify immunoreactive area from thresholded images (default method, black color, HSB color space = hue (0/29), saturation (109/253), brightness (0/220) with particle size of 600-infinity and circularity of 0.11-1.0).

### Statistics

All statistical analyses were conducted using SPSS (IBM SPSS Statistics, Version 29.0). Behavioral and immunohistochemical data were analyzed using univariate analyses of variance (ANOVA). Correlation analyses were conducted using Spearman’s rank-order correlation, as not all data was normally distributed. Across all analyses significance was set at p < 0.05.

## Results

The anatomical location of the IC targeted for analysis is shown in Fig. 1A, adapted from *The Rat Brain in Stereotaxic Coordinates* at bregma 2.04[19]. DYN A immunoperoxidase staining in the IC revealed differences between vapor- (Fig. 1C & 1E) and air-exposed groups (Fig. 1B & 1D). Alcohol self-administration (g/kg) was elevated in alcohol-dependent rats compared to non-dependent controls (Fig. 1F). A one-way ANOVA revealed a significant main effect of exposure (F(1,7) = 7.775, p = 0.027), confirming that chronic intermittent alcohol vapor exposure induced escalation of alcohol self-administration. Animals in the alcohol-dependent group emitted significantly more 22-kHz USVs during acute withdrawal compared to non-dependent controls (Fig. 1G), as revealed by a one-way ANOVA (F(1,7) = 12.664, p = 0.009). This indicates an increase in negative affective behavior associated with dependence. Alcohol-dependent animals displayed a significant increase in DYN A-like immunoreactivity within the IC compared to non-dependent controls, as indicated by the total immunoreactive area (Fig. 1H) (F(1,7) = 14.012, p = 0.007). These findings suggest that chronic alcohol exposure leads to enhanced DYN activation in this region.

**Figure 1.**
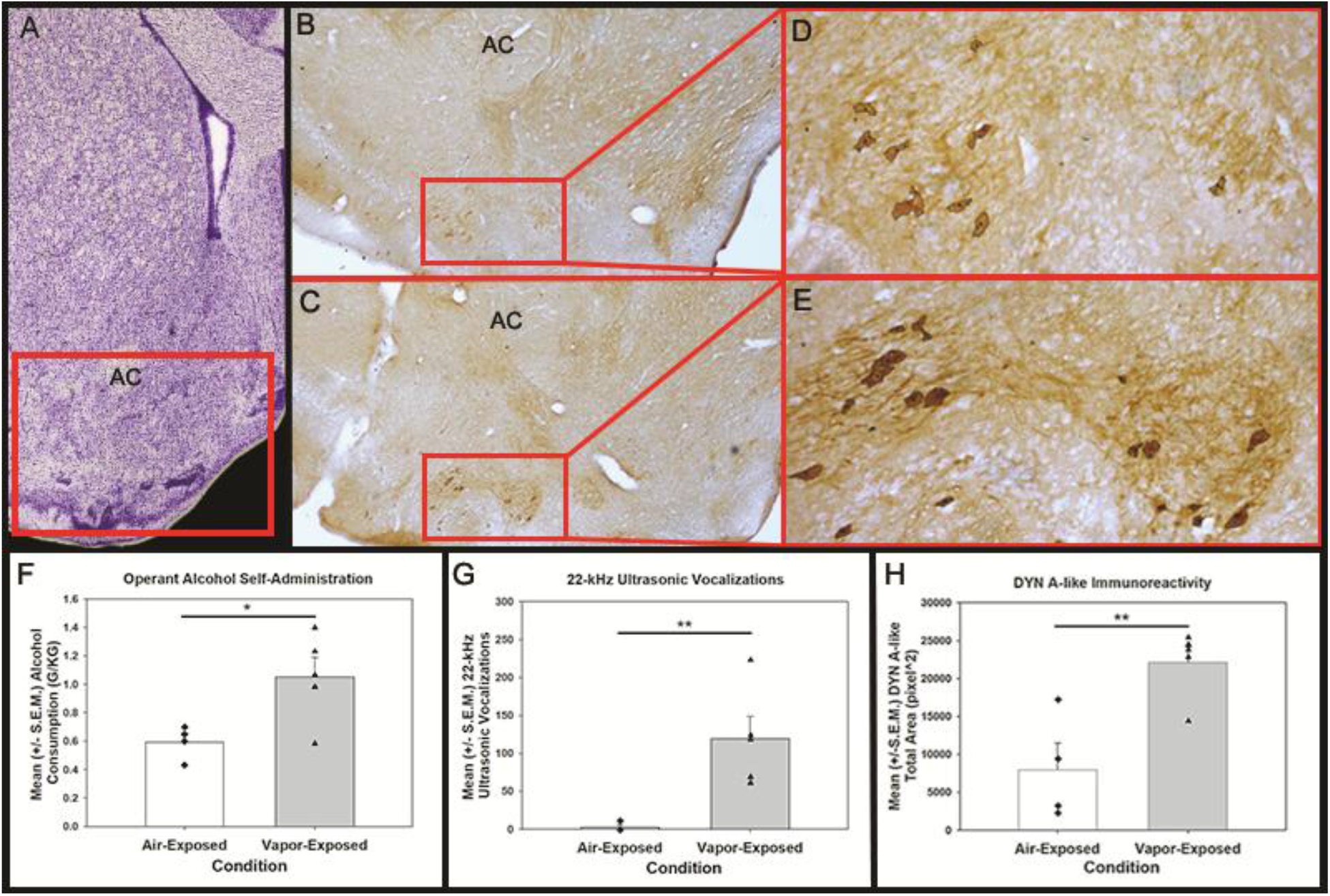
Alcohol vapor exposure increases IC DYN A immunoreactivity, alcohol intake, and USVs. The anatomical location of the IC (Fig. 1A) guided immunoperoxidase staining for DYN A in air-exposed (Fig. 1B & 1D) and alcohol vapor-exposed (Fig. 1C & 1E) rats. Vapor-exposed animals display enhanced DYN A-like immunoreactivity in the IC compared to air-exposed rats (Fig. 1H, **p < 0.01). Vapor-exposed rats demonstrate escalated alcohol self-administration (Fig. 1F, *p < 0.05). Vapor-exposed rats demonstrate a significant increase of 22-kHz USVs compared to controls (Fig. 1G, *p < 0.01).

In alcohol vapor-exposed rats, a significant positive correlation was observed between the number of 22-kHZ USVs and average DYN A neuron size in the IC (Fig. 2). Because data were not normally distributed, a nonparametric correlation analysis was performed using Spearman’s rank correlation. This analysis revealed a strong positive association between USV counts and average DYN A neuron size (ρ = 1.00, p < 0.01). In contrast, no significant correlation was observed in air-exposed animals. These findings suggest that, following alcohol vapor exposure, increased negative affective-like behavior is associated with structural alterations in DYN A-expressing neurons within the IC.

**Figure 2.**
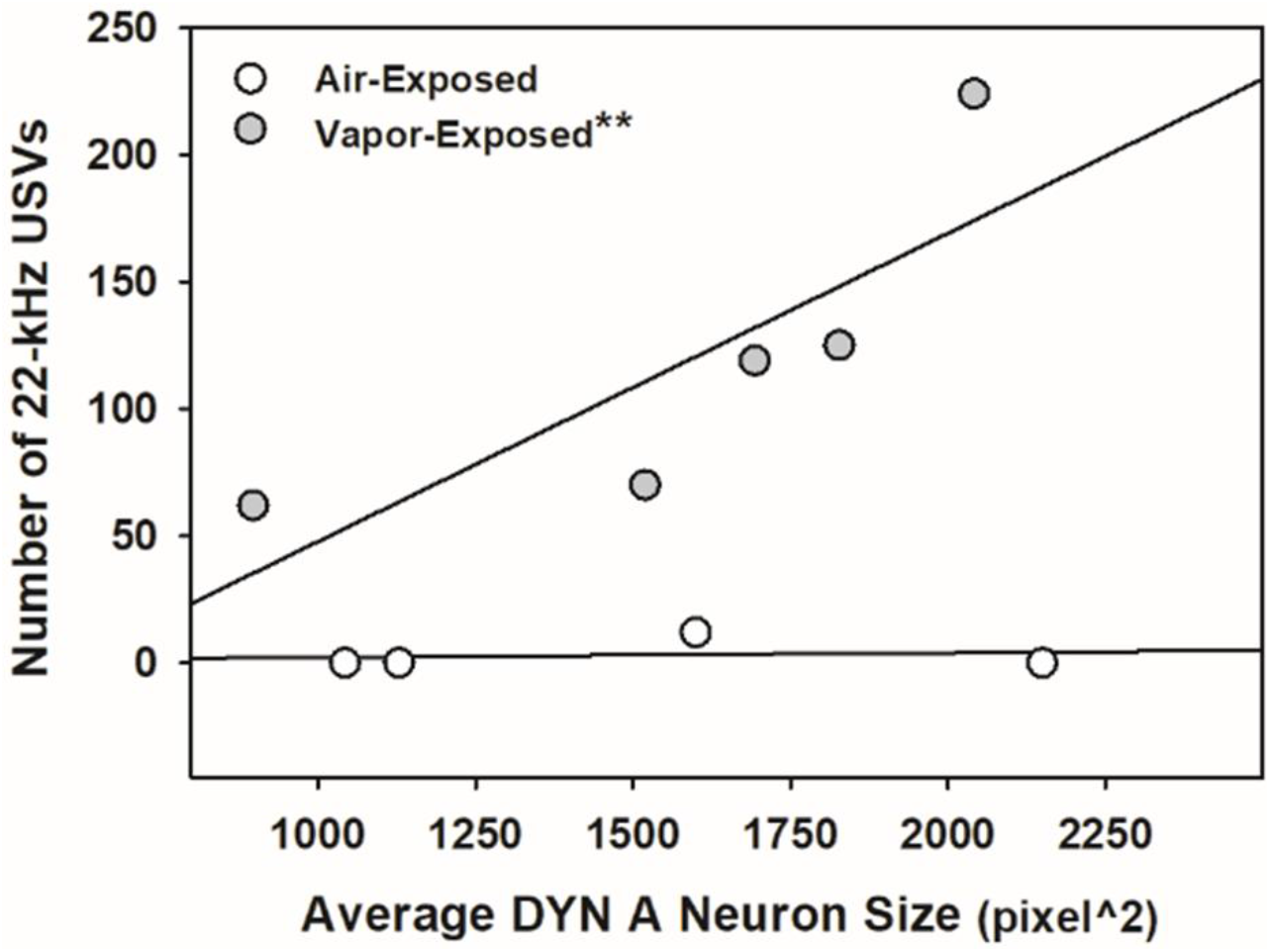
USV count correlates with IC DYN A neuron size in alcohol dependent rats. Spearman’s rank-order correlation analysis revealed a significant positive association between 22-kHz USV count and average DYN A neuron size in the IC in alcohol vapor-exposed rats (ρ = 1.00,**p < 0.01). No significant correlation was observed in air-exposed animals.

## Discussion

The present study identifies the IC as a novel locus of DYN-related neuroadaptations in the context of AUD and negative affective-like behavior. We report that CIE exposure leads to a robust escalation of alcohol self-administration (Fig. 1F) and an increase in 22-kHz USVs during acute withdrawal (Fig. 1G). These alcohol dependence-related behavioral outcomes were accompanied by significant increases in DYN A-like immunoreactivity within the IC (Fig. 1H), suggesting that this subregion of the ventral striatum may contribute to dysphoria-like states through KOR/DYN signaling.

Alcohol vapor exposed rats in the present study displayed significant negative affective behaviors that may in part be driven by disordered IC regulation. The IC receive DA input from the VTA as well as projections from other limbic and cortical structures, positioning the region at the intersection of sensory, emotional, and motivational circuits [2, 3, 20]. Robust dopamine D_3_ receptor expression has been previously established in the IC [21], and the D_3_ receptor has been previously implicated in alcohol-seeking [22]. Our findings extend this framework beyond DA interaction by demonstrating that DYN A-like immunoreactivity within the IC is associated with alcohol dependence and withdrawal-related negative affect. These results align with recent data implicating IC dysfunction in depression-like behavior and excessive grooming [6]. The observed increase in DYN A-like expression in the IC may reflect a withdrawal stress-induced upregulation of the KOR system. This is consistent with data showing that KOR activation in the VTA, NAc, and amygdala drives anxiety, dysphoria, and excessive alcohol consumption [12, 14, 15, 23]. Given the inhibitory nature of KOR signaling, increased DYN activity in the IC may suppress local DA release, resulting in reduced reward sensitivity and heightened negative affect, which could in turn drive compulsive alcohol seeking as a form of self-medication [24].

The present findings further reveal a significant positive association between 22-kHz USV count and DYN A average neuron size in the IC of alcohol dependent rats (Fig. 2). The KOR/DYN system is well established in the regulation of USVs as a marker of negative affective-like states, and prior work in our laboratory has shown that rats exhibiting elevated USVs also display increased DYN A-like immunoreactivity in the central amygdala [12]. The present results extend this relationship to the IC, suggesting that IC-localized DYN signaling could contribute to affective dysregulation associated with AUD. Furthermore, it could be that the DYN-positive neurons of the IC provide an additional source of DYN to limbic structures as has been established, for example, with medium spiny GABAergic neurons in the accumbens [25] following alcohol exposure that may contribute to states of negative affect and dysphoria.

Although the present study offers novel insight into the interplay of the IC and KOR/DYN system, certain limitations are present. While the increased DYN immunoreactivity suggests heightened DYN activity in the IC, the current study does not establish causality between IC DYN activity and behavioral outcomes. Future studies incorporating local pharmacological blockade or genetic knockdown of KOR/DYN signaling in the IC are needed to directly assess its functional relevance to withdrawal-induced negative affect and alcohol consumption. Furthermore, as males were used, we cannot rule out the potential for sex differences that remain to be established in this context. Finally, the extent to which IC DYN signaling interacts with other brain regions and stress/reward circuits remains an open and compelling question.

Importantly, these data identify the IC as a putative contributor to the affective pathology of AUD. The anatomical proximity and connectivity of the IC to limbic regions suggest a potential role in integration and expression of stress and reward signals. Our data support an expanded view of the IC as a functional contributor to affect and motivational regulation following chronic alcohol exposure. Taken together, our findings suggest the IC as a novel site of DYN system modulation in AUD, supporting the hypothesis that KOR/DYN signaling within this structure contributes to affective dysregulation during withdrawal. In highlighting the potential functional significance of the IC in AUD, this work identifies novel outcomes regarding the neurocircuitry of negative reinforcement in AUD and sets a foundation for evaluating IC-specific DYN modulation as a therapeutic strategy for AUD and other neuropsychiatric disorders.

## Acknowledgements

The authors would like to thank the USF Department of Psychiatry and Behavioral Neurosciences and the WSU Psychology Department as well as members in the Laboratory of Alcohol and Addiction Neuroscience, such as Jessica Kissler, for their assistance and the vivarium staff for their continued support.

## Author Contributions

BMW and DJR planned and completed the data collection and image acquisition and analysis. AMC, GL, GW and BMW contributed to additional data analysis and composition / editing of the manuscript.

